# A Systematic Assessment Of Current Genome-Scale Metabolic Reconstruction Tools

**DOI:** 10.1101/558411

**Authors:** S. N. Mendoza, B. G Olivier, D Molenaar, B Teusink

## Abstract

Several genome-scale metabolic reconstruction software platforms have been developed and are being continuously updated. These tools have been widely applied to reconstruct metabolic models for hundreds of microorganisms ranging from important human pathogens to species of industrial relevance. However, these platforms, as yet, have not been systematically evaluated with respect to software quality, best potential uses and intrinsic capacity to generate high-quality, genome-scale metabolic models. It is therefore unclear for potential users which tool best fits the purpose of their research. In this work, we performed a systematic assessment of the current genome-scale reconstruction software platforms. To meet our goal, we first defined a list of features for assessing software quality related to genome-scale reconstruction, which we expect to be useful for the potential users of these tools. Subsequently, we used the feature list to evaluate the performance of each tool. In order to assess the similarity of the draft reconstructions to high-quality models, we compared each tool’s output networks with that of the high-quality, manually curated, models of *Lactobacillus plantarum* and *Bordetella pertussis*, representatives of gram-positive and gram-negative bacteria, respectively. We showed that none of the tools outperforms the others in all the defined features and that model builders should carefully choose a tool (or combinations of tools) depending on the intended use of the metabolic model.

**Author Summary:** Metabolic networks that comprise biochemical reactions at genome-scale have become very useful to study and predict the phenotype of important microorganisms. Several software platforms exist to build these metabolic networks. Based on different approaches and utilizing a variety of databases it is, unfortunately, unclear what are the best scenarios to use each of these tools. Hence, to understand the potential uses of these tools, we created a list of relevant features for metabolic reconstruction and we evaluated the tools in all these categories. Here, we show that none of the tools is better than the other in all the evaluated categories; instead, each tool is more suitable for particular purposes. Therefore, users should carefully select the tool(s) that best fit the purpose of their research. This is the first time these tools are systematically evaluated and this overview can be used as a guide for selecting the correct tool(s) for each case.

## Introduction

Genome-scale metabolic models (GSMM’s) have been a successful tool in Systems Biology during the last decades [1,2], largely due to the wide range of areas for which the scientific community have found an application. GSMMs, for example, predict cellular behavior under different biological conditions, or can be used to design drug targets for important pathogens; they help to design improved strains through metabolic engineering strategies or to predict metabolic interactions in microbial communities; they have been used to study evolutionary processes or to give a rationale to lab experiments (see excellent reviews [3,4]).

The reconstruction process that forms the basis of a GSMM is very time consuming. Usually, this process starts with the annotation of a genome and the prediction of candidate metabolic functions at a genome-scale. The draft reconstruction is then refined by the user in an iterative manner though an exhaustive review of each reaction, metabolite and gene in the network. After curation, the genome-scale metabolic reconstruction is transformed into a mathematical structure, an objective function is given, constraints are set to account for specific media conditions and the resulting GSMM is evaluated to try to reproduce the experimental data. This iterative process of manual refinement is the limiting step of the whole process because it continues until the GSMM achieves a desired performance determined by the model builder. Hundreds of GSMMs have been reconstructed using this procedure, for which protocols have been described [5] and reviews are available [6,7].

Several genome-scale reconstruction tools have been developed over the last 15 years to assist researchers in the reconstruction process [8,9]. These tools are designed to speed up such process by automating several tasks that otherwise should be performed manually, such as draft network generation or gap-filling, and/or by providing useful information to the user to curate the reconstruction. There has been an outstanding increase in the number of new tools for genome-scale reconstruction which reflects the increasing interest to create high-quality GSMMs [10]. Consequently, there is a need for a systematic assessment of the performance of these tools, as many researchers are uncertain which tool to choose when they want to reconstruct their favorite organisms.

In this work, we installed and applied the most promising genome-scale reconstruction tools to provide a systematic evaluation of their performance and outputs. With each tool we reconstructed draft networks for *Lactobacillus plantarum* [11] and *Bordetella pertussis* [12], representatives of gram-positive and gram-negative bacteria, respectively, and for which high-quality GSMMs already exist. We used the manually curated GSMMs as a benchmark to assess the features of the tool-generated draft models.

### Current state of genome-scale reconstruction tools

Here we provide a brief description of the current reconstruction tools (see also S1 Table).

#### AutoKEGGRec (2018)

AutoKEEGRec [13] is an easy-to-use automated tool that uses the KEGG databases to create draft genome-scale models for any microorganism in that database. It runs in MATLAB and is compatible with COBRA Toolbox v3 [14]. One of the advantages of this tool is that multiple queries (microorganisms) can be processed in one run making it appropriate for cases where several microorganisms need to be reconstructed. The main limitation of this tool, which is directly related to the use of the KEGG database, is the lack of a biomass reaction, transport and exchange reactions in the draft genome-scale models.

#### AuReMe (2018)

AuReMe [15] (Automatic Reconstruction of Metabolic Models) is a workspace that ensures a good traceability of the whole reconstruction process, a feature that makes this tool unique. A Docker image is available for AuReMe, so users are easily able to run AuReMe in any platform without having to pre-install required packages (Windows, Linux or Mac). AuReMe creates GSMMs with a template-based algorithm [16] but it is also designed to incorporate information from different databases such as MetaCyc [17] and BIGG [18].

#### CarveMe (2018)

CarveMe [19] is a command-line python-based tool designed to create GSMMs, ready to use for Flux Balance Analysis (FBA), in just a few minutes. Its unique top-down approach involves the creation of models from a BIGG-based manually curated universal template. The implementation of its own gap-filling algorithm allows this tool to prioritize the incorporation into the network of reactions with higher genetic evidence. The authors of this tool showed that the performance of the generated models is similar to the manually curated models.

#### MetaDraft (2018)

MetaDraft [20,21] is a Python-based user-friendly software designed to create GSMMs from previously manually curated ones. It contains in its internal database BIGG models ready to be used as templates although any other model can be used as template. Users can define a specific order of templates in order to prioritize the incorporation of information related to reactions if there is a reaction match in two or more templates. One of the advantages of Metadraft is that it supports the latest features of the current SBML standards, i.e. SBML Level 3 [22] including the FBC Version 2 [23] and Groups packages [24].

#### RAVEN version 2 (2018)

RAVEN [25] (<u>R</u>econstruction, <u>A</u>nalysis and <u>V</u>isualization of M<u>e</u>tabolic <u>N</u>etworks) is a tool for genome-scale metabolic reconstruction and curation that runs in MATLAB is compatible with COBRA Toolbox v3 [14]. In contrast to the first version which only allowed reconstruction using the KEGG database [26], this evaluated version also allows the novo reconstruction of GSMMs using MetaCyc and from template models. Additionally, algorithms to merge network from both databases are provided inside RAVEN. The addition of MetaCyc allows the incorporation of transporters and spontaneous reactions to the reconstructed networks.

#### ModelSEED version 2.2 (2018)

ModelSEED [27] is a web resource for genome-scale reconstruction and analysis. This tool allows creation of GSMMs, not only for microorganisms but also for plants. The first step of its pipeline for genome-scale reconstruction is the genome annotation which is performed by RAST [28]. Users can select or even create a media to be used for gap-filling. In contrast to the first version, the second version allows creation of models in less than ten minutes (including annotation) and it provides aliases/synonyms of reactions and metabolites in other databases.

#### Pathway Tools version 22.0 (2018)

Pathway tools [29] is a software environment that support creation and curation of organism-specific databases. One of the most useful features is that users can interactively explore, visualize and edit different components of the created databases such as genes, operons, enzymes (including transporters), metabolites, reactions and pathways. Also, visualization of the whole network is possible by using Cellular Overview diagrams, in which experimental data such as gene expression can be mapped using different colors depending on the expression level.

#### Merlin version 3.8 (2018)

Merlin [30] is a java application for genome-scale reconstruction based on the KEGG database. One of the most useful resources of Merlin is the reannotation of genomes though the online service of BLAST (EBI) or HMMER. Several parameters in the annotation algorithms such as the expected value threshold and the maximum number of hits can be changed by the user if required, which makes this tool very flexible. The interface allows to compare gene function agreement between the annotation and UniProt providing information to the user for manual curation.

#### Kbase (2018)

Kbase [31] (The United States Department of Energy Systems Biology Knowledgebase) is an open-source software that allows, among a variety of functions, the reconstruction and analysis of microbes, plants and communities. Kbase is a platform that integrates several tasks such as annotation, reconstruction, curation and modeling, making suitable for the whole process of reconstruction. One of the unique features of this software is the use of narratives which are tutorials where users can interactively learn particular topics and reproduce previous results.

#### CoReCO (2014)

CoReCo [32] (<u>Co</u>mparative <u>Reco</u>nstruction) is a novel approach for the simultaneous reconstruction of multiple related species. The pipeline of CoReCo includes two steps: First, it finds proteins homologous to the input set of protein-coding sequences for each species. Second, it generates gapless metabolic networks for each species based on KEGG stoichiometry data. Thus, CoReCo allows a direct comparison between the reconstructed models, e.g. to study evolutionary aspects.

#### MEMOSys version 2 (2014)

MEMOSys [33] (<u>Me</u>tabolic <u>Mo</u>del Reseach and development System) is a database for storing and managing genome-scale models, rather than a reconstruction tool. This tool allows tracking of changes during the development of a particular genome-scale model. 20 genome-scale models are publicly available for exporting and modifying. Child models can be created from the 20 available models and then modified and compared with parent models. All the differences between different version of the models can be listed to track changes in the networks.

#### FAME (2012)

FAME [34] (<u>F</u>lux <u>A</u>nalysis and <u>M</u>odeling <u>E</u>nvironment) is a web-based application for creating and running GSMMs. This tool can reconstruct genome-scale models for any microorganism in KEGG database. One of the most interesting features of FAME is that analysis results can be visualized on familiar KEGG-like maps. It is foremost a tool for running and analyzing models and is used-by us-for educational purposes. One of the limitations of FAME is that models cannot be generated for microorganisms which are not in KEGG database.

#### GEMSiRV (2012)

GEMSiRV [35] (<u>Ge</u>nome-scale <u>M</u>etabolic Model S*i*mulation, <u>R</u>econstruction and <u>V</u>isualization) is a software platform for network drafting and editing. A manually curated model is used as template to generate a draft network for the species under study. Among the tools inside the toolbox, MrBac [36] can be used to generate reciprocal orthologous-gene pairs which are then used by GEMSiRV to generate the draft model. One of the limitations of this tool is that only one template can be used per run.

#### Other tools

Microbes Flux (2012) [37], Subliminal (2011) [38] and GEMSystem (2006) [39] are no longer maintained, as confirmed by the authors of the corresponding articles.

## Results

To assess the reconstruction tools, we performed both a qualitative and quantitative evaluation. As a first step, we created a list of relevant features for genome-scale reconstruction and software quality and we scored each tool depending on the performance (1: poor, 5: outstanding). These features are related to software performance, ease of use, similarity of output networks with regard to manually-curated models and adherence to common data standards. The criteria to assign a particular score in each feature is specified in S2 Table. Many of these features have not been assessed in previous reviews [8,9].

Subsequently, to assess how similar the generated draft networks are to high-quality models, we reconstructed with different reconstruction tools the metabolic networks of two bacteria for which high-quality manually-curated genome-scale models already were available. We chose to reconstruct the metabolic network of *Lactobacillus plantarum* and *Bordetella pertussis*, representatives of gram-positive and gram-negative bacteria, respectively. These microorganisms were selected because the corresponding GSMMs are not stored in the BIGG database, so tools that are able to use the BIGG database (AuReMe, CarveME, MetaDraft, RAVEN) in the reconstruction process cannot use the specific information for these microorganisms. If *E. coli* or *B. subtilis* would have been chosen instead we would have favored these tools because high-quality models for *E. coli* or *B. subtilis* already exist in the BIGG database and they would have been used as templates or inputs. The networks were generated using seven tools: AuReMe, CarveMe, Merlin, MetaDraft, ModelSEED, Pathway Tools and RAVEN. These cover most of the freely available software platforms. The general features of these tools are listed in Table 1.

**Table 1.**
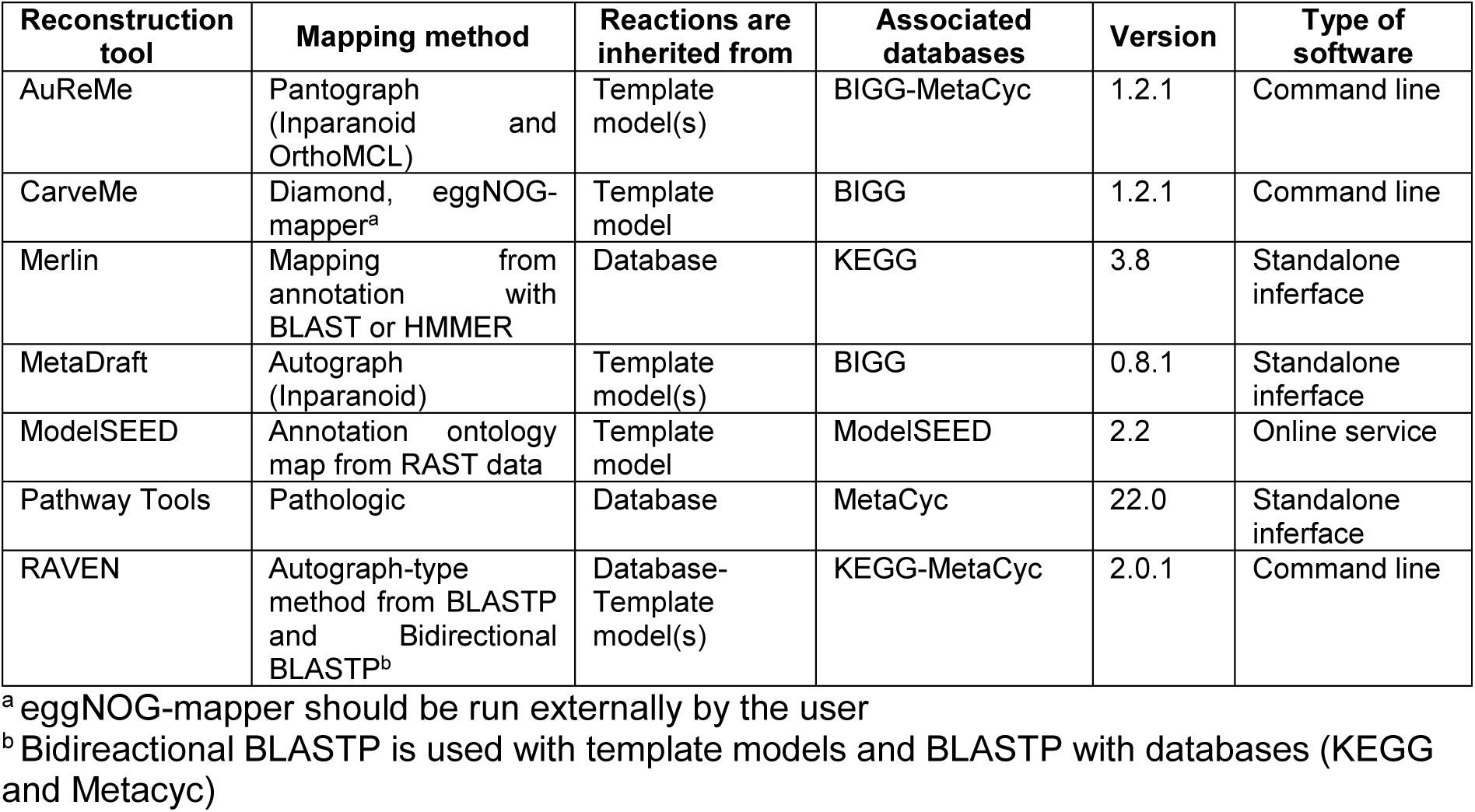
List of selected genome-scale metabolic reconstruction tools and their main features.

### General assessment overview

None of the tools got a perfect score for all of the evaluated features and usually, strengths in some tools are weaknesses in others (Fig 1, S3 table to see detailed evaluation). For example, on the one hand, ModelSEED and CarveMe were evaluated as outstanding when we checked whether the whole reconstruction process is automatic; Merlin was evaluated as poor because users should interfere more to get a network ready to perform FBA. On the other hand, we consider Merlin as outstanding with respect to a workspace for manual refinement and information to assist users during this step; CarveMe and ModelSEED do not provide further information for manual refinement nor a workspace for manual curation, so they were evaluated as poor in this category.

**Fig 1.**
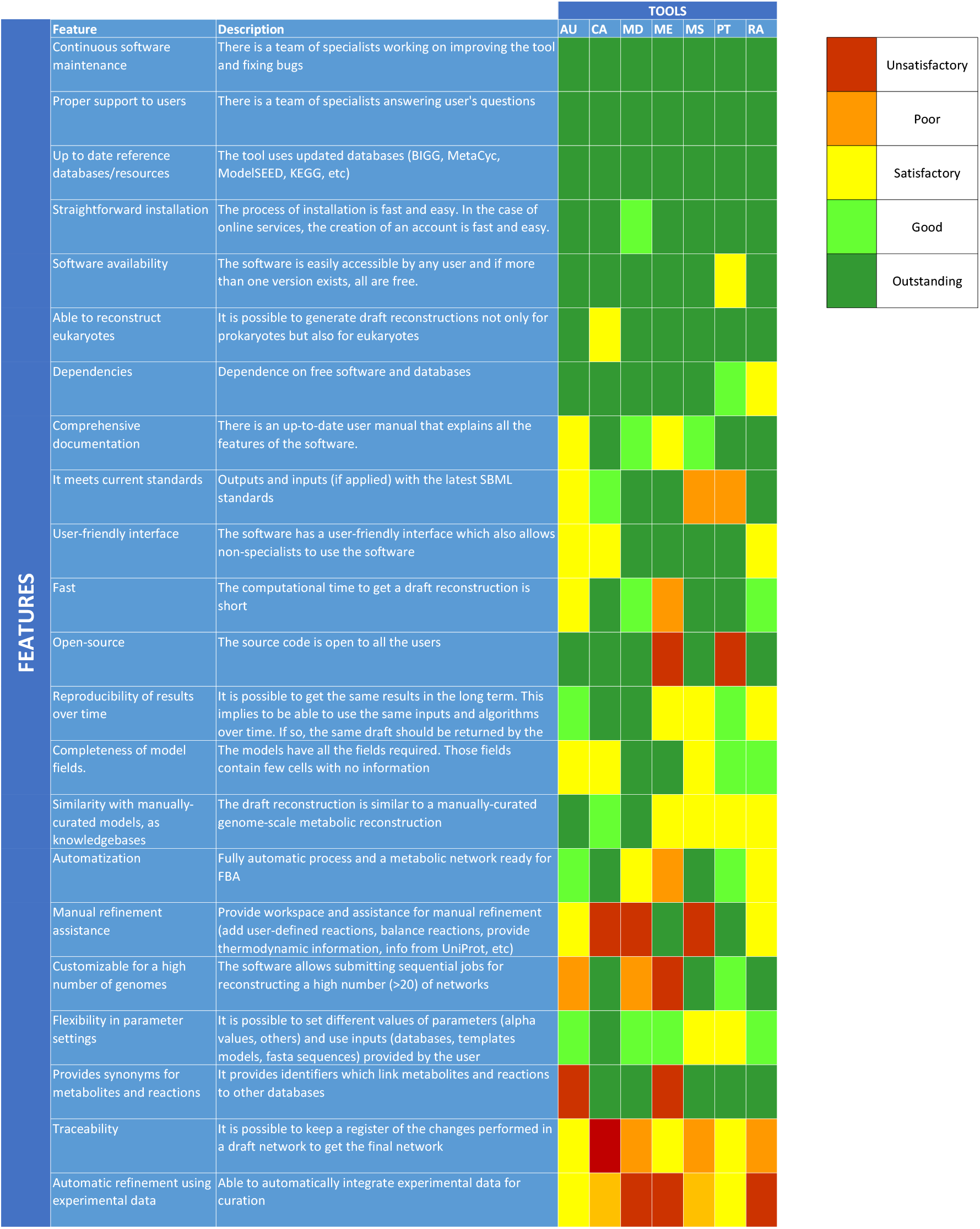
List of important features for genome-scale reconstruction and software quality and qualitative assessment of the studied genome-scale metabolic reconstruction tools. We evaluated each of the tools (AU: AuReeMe. CA: CarveMe. MD: MetaDraft. ME: Merlin. MS: ModelSEED. PT: Pathway Tools. RA: RAVEN) from an unsatisfactory (red) to an outstanding performance (dark green). In some categories such as continuous software maintenance and proper support, on the top of the figure, all the tools got the maximum score while in others such as automatic refinement using experimental data, none of the tools got the maximum. In most of the cases, strengths in some tools are weaknesses in others

In some cases, all the tools got the maximum score possible. For instance, all the tested tools are properly supported by specialist teams and also maintain up-to-date databases. In other cases, none of the tools got the maximum score. This was the case for automatic refinement of networks using experimental data. Some of the tools, such as ModelSEED and CarveMe can use media composition to gap-fill the network. AuReMe and Pathway Tools also can use, in addition to media composition, known metabolic products to gap-fill the network. In spite of that, none of the tools can also use Biolog phenotype arrays, knockout experiments and different types of omics data (transcriptomic, proteomic, metabolomic, etc.) to automatically curate the network. Although some efforts have been done in this area [40–43], this seems like a major challenge for future tool development that should lead to improved metabolic reconstructions.

Compliance with the latest SBML standards has been pointed as one of the critical points to share and represent models [44]. Consequently, we evaluated if the tools use the latest SBML features in the import (inputs) and export (outputs) of networks. For inputs, we checked if the tools were able to read networks in SBML Level 3 [22]. We additionally checked if the output networks satisfy the following three features: use of SBML Level 3 [22] with FBC annotations [23], SBML Groups [24] and MIRIAM compliant CV annotations [22,45]. These features are used, for example, for models in the BIGG database and they ensure that the information is stored in a standard way. For inputs, we found that among the tools that are able to import and use networks (AuReMe, MetaDraft, RAVEN) MetaDraft and RAVEN are able to use SBML Level 3 while AuReMe is not. For outputs, MetaDraft and Merlin and RAVEN were the only ones that exported the networks with all the three features. However, the networks created with RAVEN only had MIRIAM fields in the MAT files and they lack MIRIAM compliant CV annotations in the SBML files. In addition, AuReMe and CarveMe lack MIRIAM compliant CV annotations and SBML Groups, and Pathway Tools and ModelSEED exported the networks in SBML Level 2.

### Network comparison

We reconstructed draft networks for both *Lactobacillus plantarum* WCFS1 and *Bordetella pertussis Tohama I* with each reconstruction tool. *L. plantarum* is a lactic acid bacterium (LAB), used in the food fermentation industry and as a probiotic [46–48]. Its GSMM comprises 771 unique reactions, 662 metabolites and 728 genes, and it has been used to design a defined media for this LAB [49], to explore interactions with other bacteria [50] and as a reference for reconstructing other LAB [51]. In contrast to this LAB, *B. pertussis* is a gram-negative bacterium, and the causative agent of the Whooping cough, a highly-contagious respiratory disease [52]. The metabolic network of this pathogen was recently reconstructed and it comprises 1672 unique reactions, 1255 metabolites and 770 genes.

In total, 29 networks were created for *L. plantarum* and 27 for *B. pertussis*. The specific inputs and parameters for creating each network can be found in S1 File. Genes, metabolites and reactions were extracted from the SBML files and compared with those in the manually curated model. For convenience, the manually curated model of *L. plantarum* will be called hereafter iLP728, and the manually curated model of *B. pertussis*, iBP1870

#### Comparison of gene sets

Genes are the basis from which genome-scale model are reconstructed. When a gene is included in a metabolic reconstruction, there is at least one biochemical reaction associated with that gene. When a gene is not in the reconstruction, either the reconstruction tool couldn’t find an orthologous gene in the reference database or an orthologous gene was found but no biochemical reaction is associated to that gene. Genes sets are interesting to compare because if a gene present in the manually-curated model is absent in a draft reconstruction, that could explain why some biochemical reactions are missing in the draft. Alternatively, if a gene is absent in the manually-curated model but present in a draft reconstruction, that could explain the presence of reactions that should not be in the reconstruction. Moreover, gene sets are simple to compare among reconstructions because gene identifiers in all the cases are the same (the locus tag in the genome annotation) and so, in contrast to metabolites and reactions, there is no mapping-related bias in the comparison.

To assess how similar the draft networks were to the corresponding manually curated networks we calculated the Jaccard Distance (*JD*) as well as the ratio between the percentage of covered genes and the percentage of additional genes (*R*) (S4-S7 Tables). The *JD* has been used before to measure distance between genome-scale metabolic reconstructions, based on reaction sets [53]; here we also applied it to compare reconstructions in terms of genes and metabolites. We called *JD_g_, JD_r_* and *JD_m_* to the *JD* between two reconstructions when they are compared in terms of genes, reactions and metabolites, respectively. Analogously, we called *R_g_, R_r_* and *R_m_* to the *R* when reconstructions are compared in terms of genes, reactions and metabolites, respectively. In general terms, a value of 0 in the *JD* means that the networks are identical and a value of 1 means that the networks are totally different. For the *R*, higher values reflect a higher similarity to the original network and lower values reflect a lower similarity with the original network.

The values in the *JD_g_* ranged from 0.38 to 0.60 in *L. plantarum* and from 0.45 to 0.67 in *B. pertussis* (S4 and S5 Tables), while values in the *R_g_* ranged from 1.19 to 13.16 in *L. plantarum* and from 0.84 to 3.54 in *B. pertussis* (S6 and S7 Tables). Although the similarity of the generated draft networks seems slightly better for *L. plantarum* than for *B. pertussis*, we found that it depends on which metric is analyzed. With the exception of one network, the *R_g_* showed that all the draft networks of *L. plantarum* were more similar to iLP728 than the draft networks of *B. pertussis* to iBP1870, using the analog parameter settings. In contrast, the *JD_g_* showed that AuReMe, ModelSEED, RAVEN and Merlin generated draft networks of *L. plantarum* which are more similar to iLP728 than the draft networks of B. pertussis with regard to iBP1870, and that CarveMe, MetaDraft and Pathway Tools generated draft networks slightly more similar for *B. pertussis*.

Additionally, when sorting the values of both metrics, we noticed that the *JD_g_* order does not correspond to that made with the *R_g_*. The lowest *JD_g_* among the draft reconstructions for *L plantarum* was obtained in the network generated with AuReMe when the gram-positive set of templates was used; for *B. pertussis*, it was obtained with MetaDraft. In contrast, the highest *R_g_* among the draft reconstructions for *L. plantarum* was obtained in the network generated with AuReMe when only *Lactococcus lactis* was used as template; for *B. pertussis*, it was obtained with MetaDraft when *Escherichia coli* template was used.

Although the similarity scores for both metrics are not entirely consistent, some trends were observed. The networks more similar, in terms of genes, to the manually curated models were generated by MetaDraft, AuReMe and RAVEN (Fig 2). However, since parameters settings and inputs have a big effect on the similarity scores, the usage of these tools does not automatically ensure obtaining a draft network similar, in terms of genes, to a manually curated model. This is particularly true for RAVEN which also generated some networks with high *JD_g_* and low *R_g_* scores.

**Fig 2.**
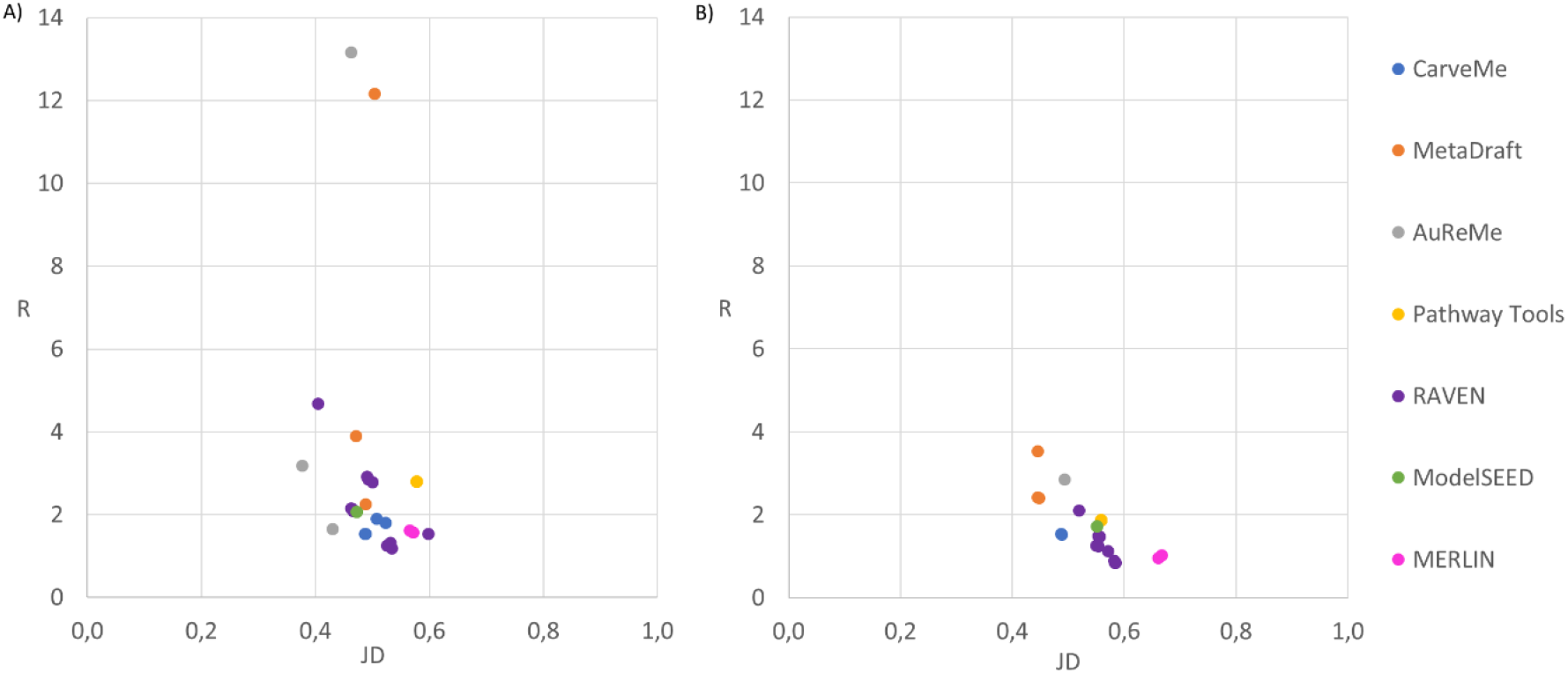
Jaccard distance versus the ratio between coverage and percentage of additional genes for draft reconstructions of *Lactobacillus plantarum* (A) and *Bordetella pertussis* (B). We used the Jaccard distance and the ratio to measure the similarity between the draft reconstructions and the corresponding manually curated models, in this case, when the networks are analyzed in terms of genes. The networks more similar to the manually curated models are located on the top left side of the plot. Thus, the draft reconstructions more similar to the manually curated models were created by AuReMe, MetaDraft and RAVEN.

We further analyzed the percentage of genes covered in the manually curated models and the percentage of genes not in the manually curated models to explain differences in *R_g_*. For both species we observed a wide variation in both variables (Figs 3 and 4). Among the five networks of *L. plantarum* with the highest coverage, two were created with AuReMe and three with RAVEN; for *B. pertussis*, four were created with RAVEN and one with CarveMe. However, the networks created with RAVEN that recovered the highest percentages of genes also added an important number of genes which were not present in the manually curated models, decreasing the values in the *R_g_*. In addition, AuReMe and MetaDraft created conservative draft networks with the lowest number of additional genes, which explains the higher values in the *R_g_*. Finally, tools such as ModelSEED, Pathway Tools and Merlin consistently created reconstructions with gene coverages not ranging in the highest values (in comparison with other networks) and adding a relatively large number of genes not present in the manually curated models, which explains why they had lower values in the *R_g_*.

**Fig 3.**
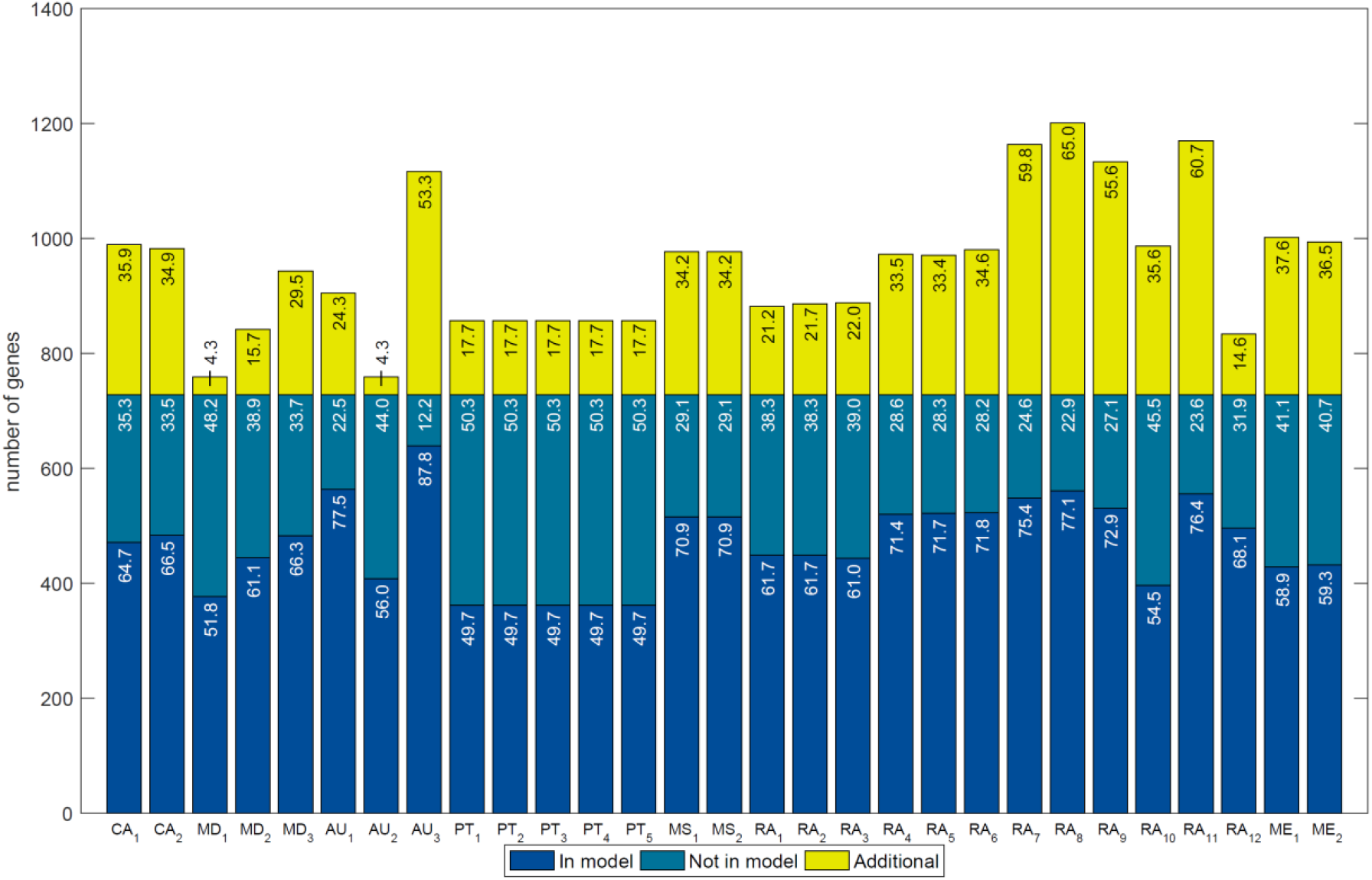
Overlap of genes in draft reconstructions for *Lactobacillus plantarum* with those in the manually-curated model, iLP728. In total, 29 networks were reconstructed with 7 tools (CarveMe: CA, MetaDraft: MD, AuReMe: AU, Pathway Tools: PT, ModelSEED: MS, RAVEN: RA, Merlin: ME). Several reconstructions, which are represented with different sub-indices, were generated for each tool using different parameters settings. Numbers inside bars represent percentages with respect to the total number of genes in iLP728. The coverage (blue bars) ranged from 49.7% to 87.8% while the percentage of additional genes (yellow bars) ranged from 4.3% to 65.0%. Most of the genes that were not recovered (dark green bars) are related to very specific metabolic functions that were carefully incorporated during the manual curation of iLP728 such as polysaccharide biosynthesis and transport.

**Fig 4.**
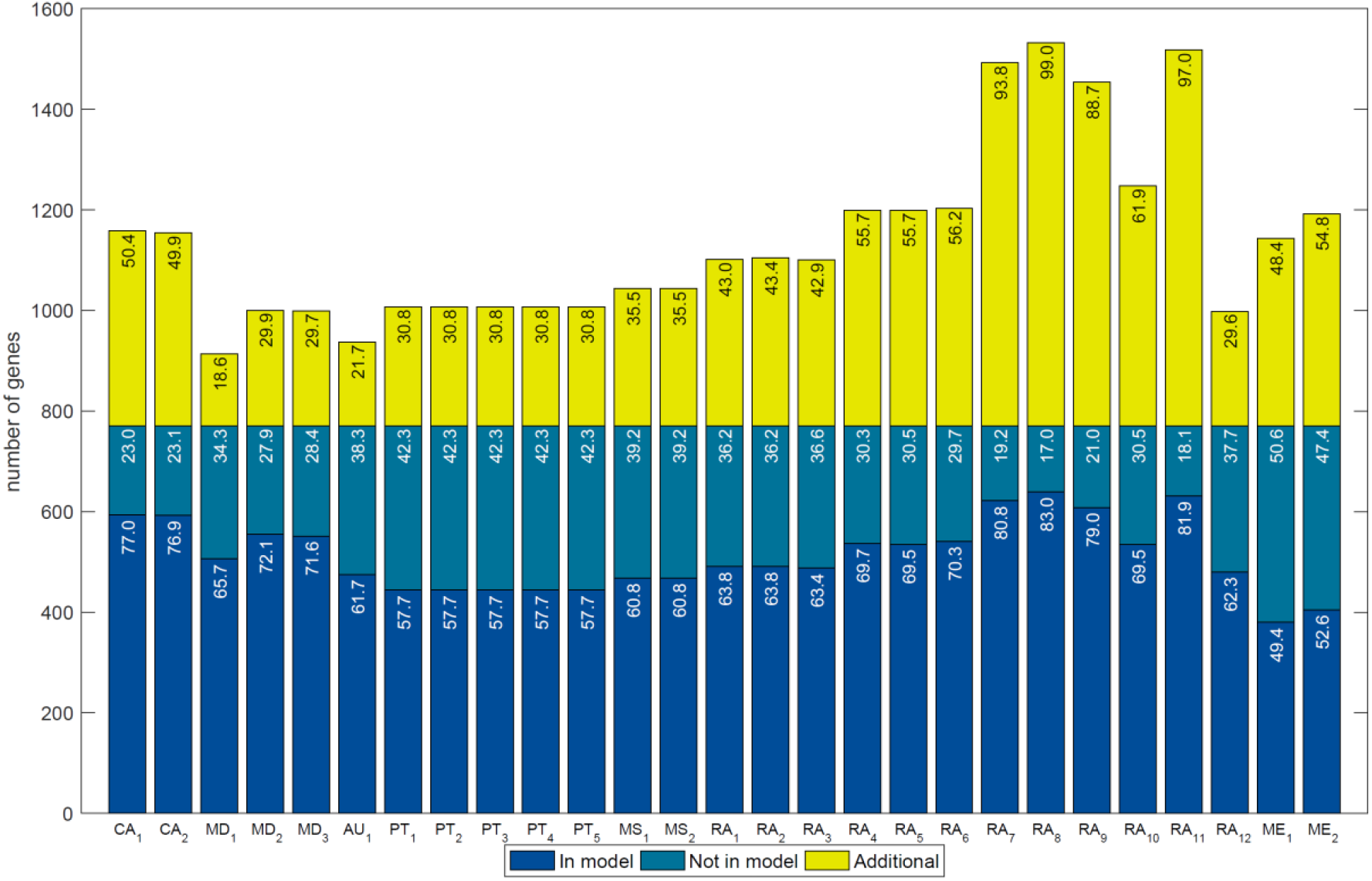
Overlap of genes in draft reconstructions for *Bordetella pertussis* with those in the manually curated model, iBP1870. In total, 27 networks were reconstructed with 7 tools (CarveMe: CA, MetaDraft: MD, AureME: AU, Pathway Tools: PT, RAVEN: RA, Merlin: ME). Several reconstructions, which are represented with different sub-indices, were generated for each tool using different parameters settings. Numbers inside bars represent percentages with respect to the total number of genes in iBO1870. The coverage (blue bars) ranged from 49.4% to 83.0% while the percentage of additional genes (yellow bars) ranged from 18.6% to 99.0%. The genes that were not recovered (dark green bars) are related to very specific metabolic functions that were carefully incorporated during the manual curation of iBP1870 such as transport and ferredoxin/thioredoxin-related reactions.

For *L. plantarum* we found 1608 different genes in total with all the tools, of which 880 were not present in iLP728. For *B. pertussis*, 1887 different genes were found, of which 1117 were not present in iBP1870. In addition, 92 genes were correctly predicted in all the draft networks for iLP728; for iBP1870 this was 121 genes. The distribution of metabolic pathways associated to those genes is wide for both species, with carbohydrate metabolism and amino acid metabolism accounting for more than 50% of the metabolic processes (S8 and S9 Tables). Additionally, 35 and 41 genes were not recovered in any network for iLP728 and iBP1870, respectively. The metabolic functions associated to those genes were very specific, with polysaccharide biosynthesis (63%) and transport (20%) top in the list for *L. plantarum* and with transport (41%) and ferredoxin/thioredoxin related reactions (26%) for *B. pertussis*. Finally, one gene in *L. plantarum*, which was associated to riboflavin biosynthesis, was recovered by all the networks but it was not present in iLP729. For *B. pertussis*, three such genes were found. These genes were associated to alternate carbon metabolism and cell envelope biosynthesis.

#### Comparison of reaction sets

Genes and biochemical reactions are connected within a reconstruction though gene-protein-reaction (GPR) associations. However, genes and reactions relationships are ultimately represented in reconstructions as boolean rules known as gene-reaction rules. With the exception of exchange, sink, demand, spontaneous and some transport reactions (e.g. those governed by diffusion), each reaction has a defined gene-reaction rule in the reference database used by each reconstruction tool. During the process of reconstruction, if orthologous genes are found that satisfy the gene-reaction rule of a particular reaction, that reaction is included in the draft reconstruction. Other reactions may be added to the draft reconstruction based on others criteria, such the probability of a particular pathway to exist in the microorganism under study or the need to fill particular gaps in the network in order to produce biomass. Nonetheless we expect that networks which are more similar in terms of genes will also be more similar in terms of reactions.

In contrast to genes, however, reactions are labeled with different identifiers in different databases. Thus, the same reaction can be stored with two different identifiers in two different databases. During the reconstruction process, reactions are added from the reference database to the draft reconstruction and tools using different databases will generate reconstructions comprising reactions with different identifiers. We therefore used MetaNetX [54] to map reactions among reconstructions built with different databases. In this approach, reactions were compared using their identifiers (case sensitive string comparison). In addition, we compared networks using reaction equations, i.e., we compared reactions using their attributes instead of their identifiers. In this second approach, we considered that two reactions were the same if they had the same metabolites with the same stoichiometric coefficients. Some exceptions were made to also match reactions that differ only in proton stoichiometry (due to differences in metabolites charge) or to catch reactions which are written in the opposite direction (reactants in the side of products).

For most networks, comparison through reaction identifiers resulted in a lower percentage of coverage than through reaction equation comparison (S10 and S11 Tables). This lower coverage was due to some missing relationships between different databases in MetaNetX, which we discovered when comparing with the reaction equations. In total, 264 new unique reaction synonyms pairs were automatically discovered for both species with the second approach (S12 Table). To further overcome the missing relationships in MetaNetX, a semiautomatic algorithm was developed to assist the discovery of new metabolite synonyms. In total, 174 new metabolites synonyms were discovered (S13 Table) which led to the discovery of 319 additional reaction synonyms (S14 Table).

The comparison through reaction equations showed a wide variation in reaction coverage and percentage of additional reactions for both species (Figs 5 and 6). In addition, for those networks created with RAVEN (KEGG), ModelSEED and Merlin we observed a large number of reactions with a partial match with the manually curated model. These partial matches emerge from differences in proton stoichiometry, which indicates the existence of an important number of metabolites with different charge than those found in the manually curated models. In contrast to the gene sets comparison, where the coverage was as high as 88% and 83%, we only observed a maximum coverage of 72% and 57%, for *L. plantarum* and *B. pertussis*, respectively, even when considering partial matches. For both species, on average, around 50% of the reactions that were not recovered in the manually-curated models don’t have gene-reaction associations. If we take all the unique genes associated to the remaining 50% of the reactions, on average, around 40% of those are not in the draft models, which can explain why we observed an important decrease in the coverage of reactions.

**Fig 5.**
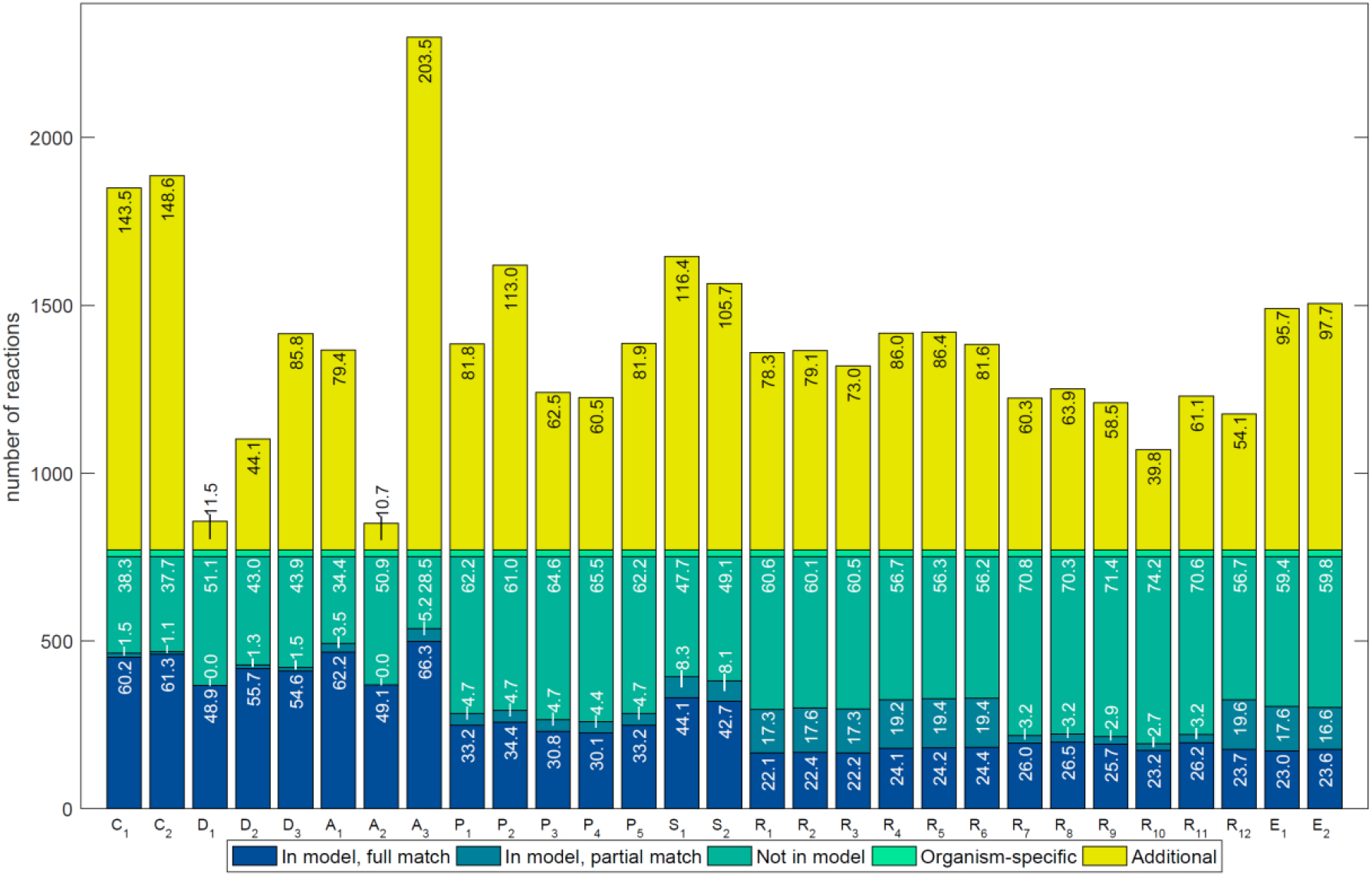
Overlap of reactions in draft reconstructions for Lactobacillus plantarum with those in the manually-curated model, iLP728. In total, 29 networks were reconstructed with 7 tools (CarveMe: CA, MetaDraft: MD, AuReMe: AU, Pathway Tools: PT, ModelSEED: MS, RAVEN: RA, Merlin: ME). Several reconstructions, which are represented with different sub-indices, were generated for each tool using different parameters settings. Numbers inside bars represent percentages with respect to the corrected number of reactions in iLP728, which is the total number of reactions in iLP728 minus the biomass-related reactions (light green). We observed a wide variation in the coverage (blue bars) and percentage of additional reactions (yellow bars). In addition, an important number of reactions in the networks build with ModelSEED, RAVEN (KEGG) and Merlin contained different stoichiometry for protons than those in iLP728 (dark green bars).

**Fig 6.**
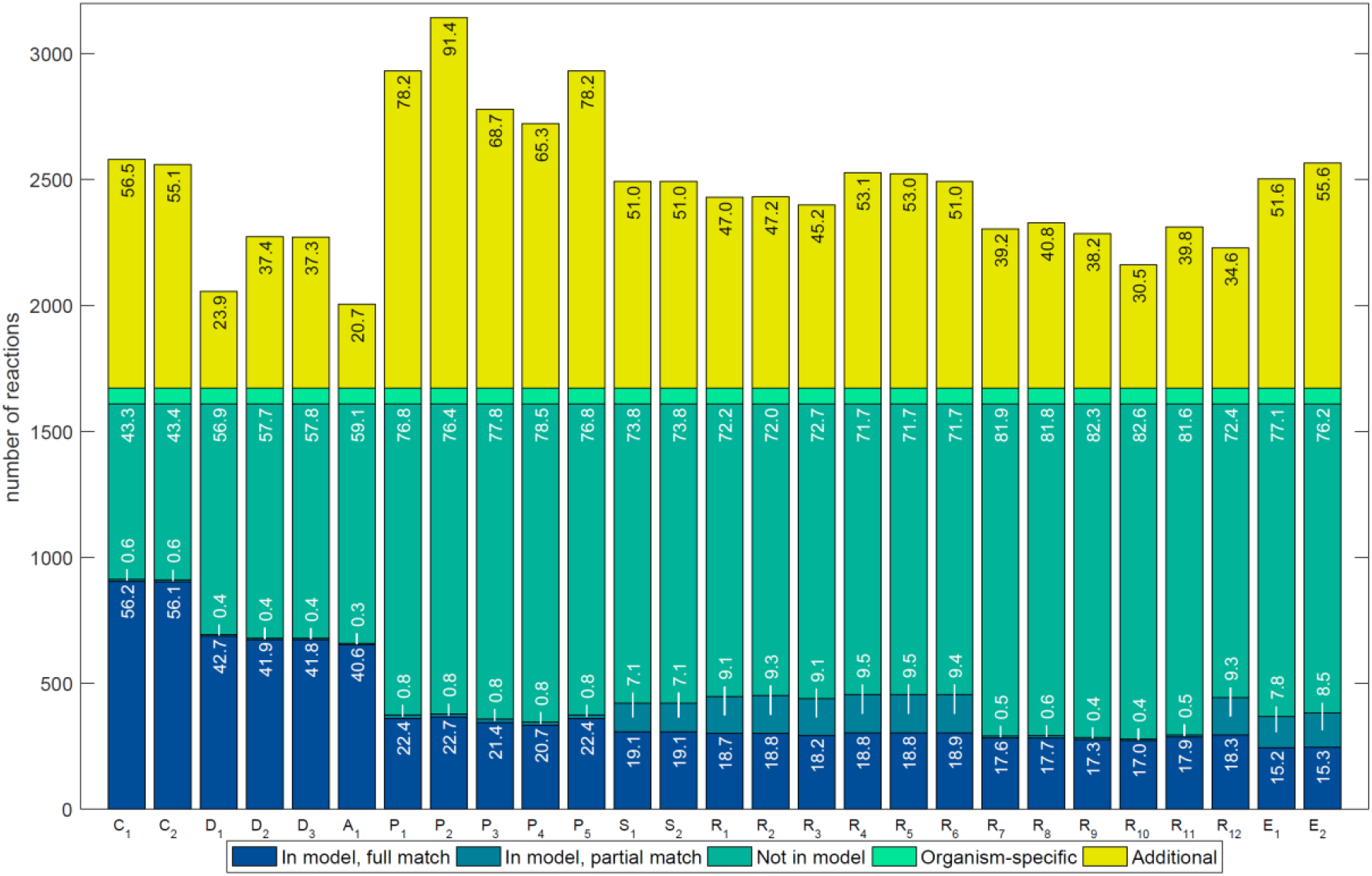
Overlap of reactions in draft reconstructions for Bordetella pertussis with those in the manually-curated model, iBP1870. In total, 27 networks were reconstructed with 7 tools (CarveMe: CA, MetaDraft: MD, AuReMe: AU, Pathway Tools: PT, ModelSEED: MS, RAVEN: RA, Merlin: ME). Several reconstructions, which are represented with different sub-indices, were generated for each tool using different parameters settings. Numbers inside bars represent percentages with respect to the corrected number of reactions in iBP1870, which is the total number of reactions minus the biomass-related reactions (light green). We observed a wide variation in the coverage (blue bars) and percentage of additional reactions (yellow bars). In addition, an important number of reactions in the networks build with MODELSEED, RAVEN (KEGG) and Merlin contained different stoichiometry for protons than those in iBP1870 (draft green bars).

We again calculated the *JD_r_* and the *R_r_* to assess how similar the networks were, in this case in terms of reactions. The first observation we made is that, independent of the metric and for both species, each reconstruction was less similar in terms of reactions than in terms of genes, which is consistent with the decrease in coverage. In addition, as in the gene comparison, the order of scores for the *R_g_* and the *R_r_* by magnitude, was not the same. If we compare the similarity scores for reactions sets with the ones for genes sets, we see almost the same trend but with one difference. AuReMe and MetaDraft are still the tools with the best similarity scores but now CarveMe goes up in the list of scores and RAVEN goes down (Fig 7, S4-S7 Tables). This was particularly true for *B. pertussis* where two networks reconstructed with CarveMe got the two first places in the *JD_r_* list.

**Fig 7.**
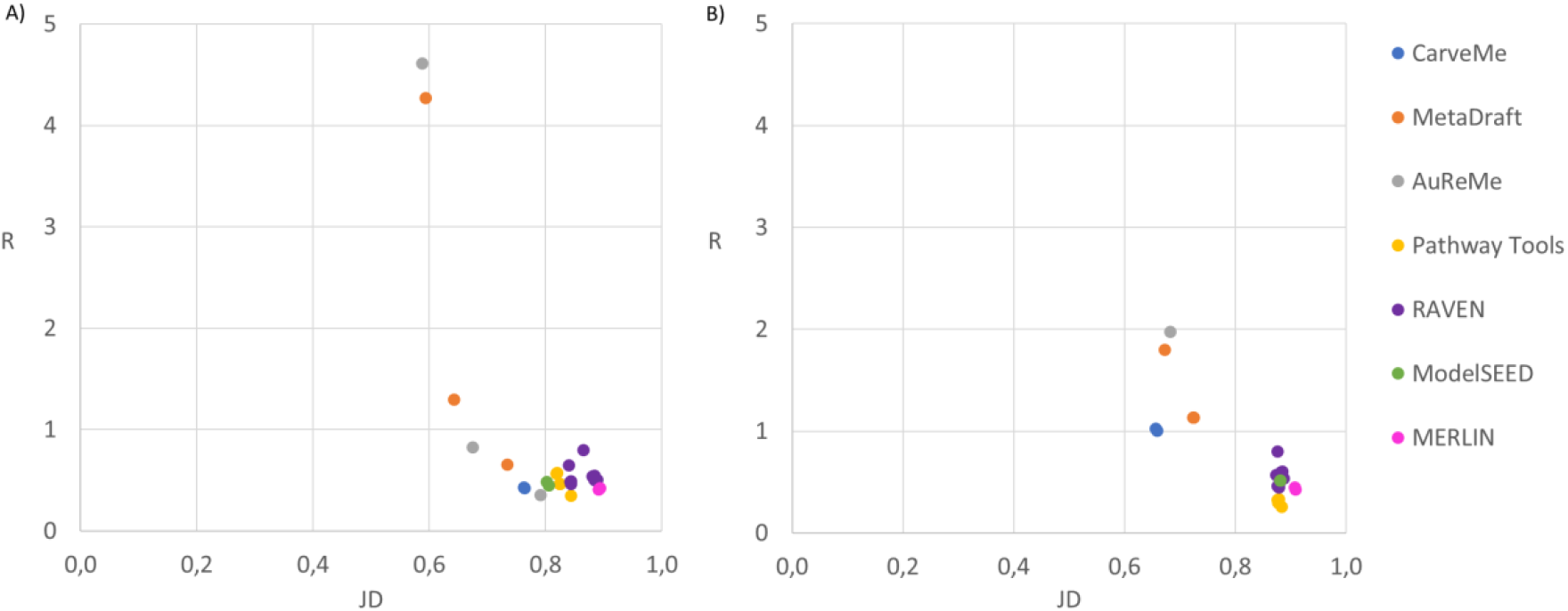
Jaccard distance versus the ratio between coverage and percentage of additional reactions for draft reconstructions of *Lactobacillus plantarum* (A) and *Bordetella pertussis* (B). We used the Jaccard distance and the ratio the measure the similarity between the draft reconstructions and the corresponding manually curated model, in this case, when the networks are analyzed in terms of reactions. The networks more similar to the manually curated models are located on the top left side of the plot. Thus, the draft reconstructions more similar, in terms of reactions, to the manually curated models were created by AuReMe, MetaDraft and CarveMe

Although RAVEN generated some reconstructions with high genes sets similarity to the manually curated models, it did not for reactions sets similarity. We therefore analyzed one of the networks reconstructed with RAVEN in more detail, one that was consistently in the top-5 list for both species for both metrics. We found two reasons for the decrease in performance. First, the analyzed network was created based on KEGG, so metabolites were not labelled as intracellular or extracellular. Hence no transport or exchange reactions were present. Although there are functions to incorporate this kind of reactions in RAVEN, that is considered as manual curation because users must specify which compounds should be transported, and we here only tested how much work would it take to transform these draft networks into high-quality reconstructions. Second, due to some not intuitive duplicated entries in the BIGG database, the translation of metabolites though MetaNetX dit not always result in a match with the manually curated model. For example, the reaction R04534 in the draft created with RAVEN was never mapped to the reaction HDER4 in iLP728 because in MetaNetX R04534 is associated with 3OAR100, a duplicated reaction of HDR4 in BIGG. It was also not recovered when comparing reaction equations because the metabolites in R04534 were mapped to those in 3OAR100, not to those in HDR4.

We further analyzed reactions that were present and absent in all the reconstructions to understand to which kind of metabolic processes they were related. 71 reactions in iLP728 and 91 in iBP1870 were always found in all the draft networks. In agreement with the genes sets analysis, the associated metabolic processes are mainly amino acid metabolism, nucleotide metabolism and carbohydrate metabolism (S15 and S16 Tables). Additionally, 168 reactions in iLP1870 and 609 in iBP1870 were not found by any tool. In both species, around 10% of those reactions were biomass-related reactions and from the rest, most of them were exchange reactions, transport reactions without gene associations and reactions in other categories that were not in the BIGG database (S17 and S18 Tables). Only one reaction, associated to amino acid metabolism, was found in all the draft networks of *L. plantarum* but not in iLP728; four reactions, associated mainly to carbohydrate metabolism, were found in all the draft networks but not in iBP1870.

#### Comparison of metabolite sets

Other important elements within metabolic reconstructions are metabolites. When a biochemical reaction is added to the draft network during the reconstruction process, all the reactants and products are added to the network too. As the draft metabolic networks were created with different tools, each of which using its own set of databases, they had different identifiers for the same metabolite. For those networks whose identifiers were different from BIGG, we again used MetaNetX and our own additional dictionary to map metabolites.

We calculated the *JD_m_* and the *R_m_* to assess the metabolite sets similarity. For almost all the draft networks in both species, the values in the *JD_m_* were between the *JD_g_* and the *JD_r_*; we found the same for the *R_m_* (S4 – S7 Tables). Again, when sorting the networks according to their metric scores, we found the same trends than for reactions sets. The first position in the lists were networks either reconstructed with MetaDraft, AureMe or CarveMe. Moreover, independently of the metric and the species, MetaDraft reconstructed 40% of the networks among those in the top-5.

208 metabolites in iLP728 and 266 in iBP1870 were correctly predicted in all the draft networks. These metabolites were in both cases mainly associated to carbohydrate metabolism and amino acid metabolism (S19 and S20 Tables). 86 metabolites in iLP728 and 312 in iBP1870 were not recovered in any network. Of those, 16 were related to the biomass of *L. plantarum* and 20 others were not in the BIGG database. For iBP1870, 47 were biomass-related and 127 others were not in the BIGG database. Finally, 9 and 13 metabolites were recovered in all the networks but they were not present in iLP728 and iBP1870, respectively. Mainly, they were associated to metabolism of cofactors and vitamins and amino acid metabolism in the case of *L. plantarum* and carbohydrate metabolism and glycan biosynthesis in the case of *B. pertussis* (S21 and S22 Tables).

#### Topological Analysis

To compare the topological features of each network, we calculated the number of dead-end metabolites, the number of orphan reactions, the number of unconnected reactions and other metrics (S23 and S24 Tables).

iLP728 has 113 dead-end metabolites while iBP1870 has 60. This is consistent with the observation that many pathways are disrupted in *L. plantarum* leading for example to well-known auxotrophies for many amino acids [49,55]. With the exception of CarveMe, all the tools generated networks with a high number of dead-end metabolites, ranging from 237 and 996, and from 379 to 976, for *L. plantarum* and *B. pertussis*, respectively. The low number of dead-end metabolites in CarveMe is caused by the use of a manually curated universal model as a template which lacks dead-end metabolites.

Without considering exchange and demand/sink reactions, 128 and 449 reactions without gene associations (called orphan reactions) were found in iLP728 and iBP1870, respectively. These reactions are mainly associated to transport reactions, amino acid metabolism and biomass formation. AuReMe and RAVEN returned metabolic networks with no orphan reactions. These tools only include reactions with genomic evidence and others lacking this support, such as exchange reactions, are not included. MetaDraft and ModelSEED returned networks with a low amount of orphan reactions, which are related in the case of MetaDraft to duplicated reactions and in the case of ModelSEED, to exchange reactions. In contrast, CarveMe, Pathway Tools and Merlin returned networks with a significantly larger number of orphan reactions (ranging from 66 to 345 in *L. plantarum* and from 115 to 736 in *B. pertussis*). For CarveMe, this is due to the inclusion of transport and spontaneous reactions as well as reactions needed to create biomass (from gap-filling); for Pathway tools it is because of the addition of reactions to complete probable pathways and spontaneous reactions; for Merlin, this is solely due to spontaneous reactions.

## Discussion

In this work, we reviewed the current state of all the reconstruction tools we could find in the literature and performed a systematic evaluation of seven of them. None of the tools performed well in all the evaluated categories so users should carefully select the tool(s) that suit the purpose of their investigation. For example, if a high-quality draft is required and models are available for a phylogenetically close species, MetaDraft or AuReMe could be selected, reducing thus the time needed to obtain a high-quality manually-curated model. Of these, MetaDraft was the most robust for handling models and since it has a graphical user interface, it is also suitable for non-specialists. AuReMe, on the contrary, offered a command-line workspace where the traceability is the priority. Although we were not able to use RAVEN in the template mode, this tool allowed us to automate the generation of several reconstructions, it had a high flexibility with parameters and it offered integration with the KEGG and MetaCyc databases which makes it very appropriate for less studied species. ModelSEED, CarveMe and Pathway Tools were the fastest tools to generate reconstructions having a great potential for large-scale studies how is has been proven in previous works [53,56]. The first two tools provided networks which are ready to perform FBA, however presumably because of the automatic gap-filling procedure, too many reactions that should be manually verified must be expected. Pathway Tools and Merlin provided a platform suitable for manual curation which nicely guide the user though the whole reconstruction process.

The list of features that we defined not only can be used by model builders to select the best tool(s) but also by developers as a guide for improving them. We highlight four features, which are in accordance with the FAIR guiding principles for scientific data management and stewardship [57], that should be considered as a priority by developers to ensure management of reconstructions in a standard way: 1) To be findable: All the genes, metabolites and reaction in a reconstruction should be assigned with unique and persistent identifiers, and synonyms or aliases in other databases should be provided whenever possible 2) To be accessible: exhaustive control of versions should be implemented so users will be able to submit small but significant changes to draft reconstructions, to trace changes made during the reconstruction process, or to retrieve a particular version if desired. 3) To be interoperable: output (and input if applied) reconstructions should be written with the latest features of the SBML standards 4) To be reusable: In relationship with providing a detailed provenance, transparency of decisions through the whole reconstruction process should be ensured so users can see why a particular reaction was added and at which stage (draft network generation, gap-filling, refinement, etc.).

Genome-scale reconstructions are usually evaluated after they are converted into genome-scale models [5] i.e., mathematical structures where simulations can be performed under constraints that describe specific experimental conditions. Thus, GSMMs are tested by their accuracy to predict experimental data such as knock-outs, nutritional requirements and growth rates on different conditions. However, most of the drafts we generated were not suitable to perform FBA, mainly due to the lack of biomass-related, transport and exchange reactions. Thus, we limited the evaluation of the drafts to the comparison with manually-curated, genome-scale reconstructions. The later are valuable by themselves as knowledgebases because they contain extensive information from literature. Here, we prescribed that the manually-curated reconstructions are the gold standard, which implies that they cannot be improved and that is obviously not true. Many reconstructions of for example, *E. coli, S. cerevisiae* and *H. sapiens* have gone through multiple rounds of improvements during the years [58–60]. As reference databases used by reconstruction tools increase in size and quality, so will the reconstructions which are based on them. Therefore, some of the reactions which were suggested by the tools and which are not in the manually-curated models could indeed be reactions which would improve the quality of the reconstructions. Whether one of those reactions should be in the reconstruction or not will depend not only on the genomic evidence but also on the scope and context of the reconstruction. Many reactions are usually not incorporated because they are not needed for modeling purposes [5]. Thus, similarity scores shouldn’t be taken alone to assess the quality of draft reconstructions. Indeed, additional reconstructions of *Lactobacillus plantarum* that we made with CarveMe and ModelSEED and which were gap-filled using a modified version of CDM (S2 File), a media that support the growth of this microorganism in vivo [49], showed a general performance close to the manually curated model, suggesting that although the networks are not so similar as others created with different tools, the core metabolism remains similar. Despite that, the performance of these networks is dependent on the media composition which is used for the gap-filling (S1 Fig), and therefore if there is no experimentally determined media, some false positive and false negative predictions could emerge. For example, if very accurate predictions regarding nutritional requirement are needed to design a microbial community, automatic reconstructions for which an experimentally determined media composition is not provided during gap-filling could result in false predictions.

A correct mapping of identifiers among different databases is crucial to perform a proper comparison between metabolic networks. Important efforts such as MetaNetX [54] and Borgifier [61] have been done to facilitate this titanic task. The first of those tools allowed us to map most of the metabolites and reactions among the different reconstructions but naturally some relationships were missing. To overcome this limitation, we implemented an algorithm to search reaction equations, even when they have differences in proton stoichiometry due to different protonation states or even if the reactions are written in the opposite direction. As a second step to further reduce the fraction of metabolites which were not mapped and though a semi-automatic and iterative process, we determined 174 new relationships. In spite of our efforts, still some relationships were missing which evidence the complexity of the problem. Since recent efforts have made clearer the type of issues arising in different databases [62], we emphasize the importance of standards, which could make easier the identification of synonyms because of the presence of high-quality information, and the need of an outstanding mapping system.

Systematic assessments of tools for systems biology have become very popular [63,64] due to the great impact they have in the community of potential users who certainly are searching the best tool to apply in their research. Knowing the strengths and limitations of each tool allow users to select the best tool(s) for their case, to save time in preliminary tests and to focus more on the analysis and modelling using those reconstructions. Moreover, to provide genome-scale models of high quality, in terms of usability and standards, has become a priority during the last years. Efforts such as those done by Memote [44] highlight the need for suites that test the quality of genome-scale models to ensure high-quality outputs, not only in terms of their content as knowledgebases but also in terms of standards.

## Conclusions

All the assessed reconstruction tools showed strengths and weaknesses in different areas and none of the tools outperformed the others in all the categories. In particular, template-based reconstruction tools such as AuReMe, MetaDraft and CarveMe generated networks with a higher reactions sets similarity to manually curated networks than other tools. In addition, tools such as Pathway Tools and Merlin provide a proper workspace and useful information for manual refinement which could be suited for cases where much time can be dedicated to this step. RAVEN provides a platform in which biochemical information from different databases and approaches can be merged, which could be useful for less characterized species. Finally, tools such as CarveMe and ModelSEED provide ready-to-use metabolic networks which can be useful for a fast generation of model-driven hypothesis and exploration but users will have to be aware of potential false results.

There seems to be a trade-off between coverage and similarity, and it remains to be seen how much room for improvement there is. We see three clear features that would improve any tool: better standards that would allow easier integration of the best of tools, exhaustive version control during the reconstruction process, and algorithms that can use experimental data for inclusion of genes and reactions into the models.

## Materials and Methods

### Protein sequences

We used the protein sequences or the GenBank files of the different microorganisms as input to generate the genome-scale metabolic reconstructions with each of the selected tools. All the protein sequences were downloaded from NCBI. For *Lactobacillus plantarum* strain WCFS1 and *Bordetella Pertussis* strain Tohama I we used the protein sequences deposited under the NCBI accession numbers NC_004567.2 and NC_002929.2 respectively.

### Reconstruction

The specific parameters and inputs used to reconstruct the draft networks with each tool can be found in S1 file.

#### AuReMe

We used AuReMe version 1.2.1, which was downloaded using Docker Toolbox, to generate the draft reconstructions.

To generate the genome-scale metabolic reconstructions of *Lactobacillus plantarum* we used three different set of templates from the BIGG database: 1) *Lactococcus lactis* (iNF517). 2) *Lactococcus lactis* (iNF517), *Bacillus subtilis* (iYO844), *Staphylococcus aureus* (iSB619), *Clostridium ljungdahlii* (iHN637) and *Mycobacterium Tuberculosis* (iNJ661). 3) *Lactococcus lactis* (iNF517), *Bacillus subtilis* (iYO844), *Staphylococcus aureus* (iSB619), *Clostridium ljungdahlii* (iHN637), *Mycobacterium Tuberculosis* (iNJ661), *Escherichia coli* (iML1515), *Klebsiella pneumoniae* (iYL1228), *Shigella flexneri* (iSFxv_1172), *Shigella boydii* (iSbBs512_1146), *Shigella sonnei* (iSSON_1240), *Pseudomonas putida* (iJN746), *Yersinia pestis* (iPC815), *Helicobacter pylori* (iIT341), *Geobacter metallireducens* (iAF987), *Salmonella entérica* (STM_v1_0), *Thermotoga marítima* (iLJ478), *Synechocystis sp* (iJN678) and *Synechococcus elongatus* (iJB785)

For *Bordetella pertussis* we used *Escherichia Coli* as template (iML1515).

#### CarveMe

We used CarveMe version 1.2.1 (downloaded from https://github.com/cdanielmachado/carveme on August 1^st^, 2018) to generate the draft reconstructions. Two genome-scale metabolic reconstructions were generated for *Lactobacillus plantarum* using the universal bacterial template and the gram-positive bacterial template, respectively. For *B. pertussis*, the universal bacterial template and the gram-negative bacterial template were used.

#### Merlin

We used Merlin version 3.8 (downloaded from https://merlin-sysbio.org/index.php/Downloads on August 1^st^, 2018) to generate the draft reconstructions. For all the networks, we first annotated the genomes with EBI through MERLIN using default parameters. Then, we loaded KEGG metabolic data and integrated the annotation with the model. Finally, we created gene-reaction-protein associations and removed unbalanced reactions to be able to export the network to SBML format.

#### MetaDraft

We used MetaDraft version 0.8.1, which was obtained by request from Brett Olivier on May 1^st^ of 2018, to generate the draft reconstructions. Now available from https://systemsbioinformatics.github.io/cbmpy-metadraft/.

To generate the genome-scale metabolic reconstructions of *Lactobacillus plantarum* we used three different set of templates from the BIGG database: 1) *Lactococcus lactis* (iNF517). 2) *Lactococcus lactis* (iNF517), *Bacillus subtilis* (iYO844), *Staphylococcus aureus* (iSB619), *Clostridium ljungdahlii* (iHN637) and *Mycobacterium Tuberculosis* (iNJ661). 3) *Lactococcus lactis* (iNF517), *Bacillus subtilis* (iYO844), *Staphylococcus aureus* (iSB619), *Clostridium ljungdahlii* (iHN637), *Mycobacterium Tuberculosis* (iNJ661), *Escherichia coli* (iML1515), *Klebsiella pneumoniae* (iYL1228), *Shigella flexneri* (iSFxv_1172), *Shigella boydii* (iSbBs512_1146), *Shigella sonnei* (iSSON_1240), *Pseudomonas putida* (iJN746), *Yersinia pestis* (iPC815), *Helicobacter pylori* (iIT341), *Geobacter metallireducens* (iAF987), *Salmonella entérica* (STM_v1_0), *Thermotoga marítima* (iLJ478), *Synechocystis sp* (iJN678) and *Synechococcus elongatus* (iJB785)

To generate the genome-scale metabolic reconstructions of *Bordetella pertussis* we used three different set of templates from the BIGG database: 1) *Escherichia coli* (iML1515). 2) *Escherichia coli* (iML1515), *Klebsiella pneumoniae* (iYL1228), *Shigella flexneri* (iSFxv_1172), *Shigella boydii* (iSbBs512_1146), *Shigella sonnei* (iSSON_1240), *Pseudomonas putida* (iJN746), *Yersinia pestis* (iPC815), *Helicobacter pylori* (iIT341), *Geobacter metallireducens* (iAF987), *Salmonella entérica* (STM_v1_0), *Thermotoga marítima* (iLJ478), *Synechocystis sp* (iJN678) and *Synechococcus elongatus* (iJB785). 3) *Escherichia coli* (iML1515), *Klebsiella pneumoniae* (iYL1228), *Shigella flexneri* (iSFxv_1172), *Shigella boydii* (iSbBs512_1146), *Shigella sonnei* (iSSON_1240), *Pseudomonas putida* (iJN746), *Yersinia pestis* (iPC815), *Helicobacter pylori* (iIT341), *Geobacter metallireducens* (iAF987), *Salmonella entérica* (STM_v1_0), *Thermotoga marítima* (iLJ478), *Synechocystis sp* (iJN678), *Synechococcus elongatus* (iJB785), *Lactococcus lactis* (iNF517), *Bacillus subtilis* (iYO844), *Staphylococcus aureus* (iSB619), *Clostridium ljungdahlii* (iHN637) and *Mycobacterium Tuberculosis* (iNJ661).

#### ModelSEED

We used ModelSEED version 2.2 web service on August 16^st^ of 2018 to generate the draft reconstructions. Models were created using different template models. No media was specified to create the models.

#### Pathway Tools

We used Pathway Tools version 22.0 to generate the draft reconstructions. Four networks were created with the Desktop mode using different cut-off values for pathways prediction and one was made with the Lisp-console with default parameters. All the networks were exported manually with the Desktop mode

#### RAVEN

We used RAVEN version 2.0.1, which was downloaded from https://github.com/SysBioChalmers/RAVEN, to generate the draft reconstructions. Different models were created using different databases (KEGG and MetaCyc) and different values in the parameters for orthology searches.

### Pre-processing of *L. plantarum* and *B. pertussis* network

We pre-processed the manually curated networks in order to compare them with the draft networks. We semi-automatically changed metabolite and reaction identifiers to match those of the BIGG database. Also, we removed duplicated reactions (those with the same reaction equation). Before deletion of a duplicated reaction, the associated gene-reaction rule was transferred to or merged with the gene-reaction rule of the reaction that was kept in the network.

### Comparison of gene sets

We define the union of all the unique genes found in a particular metabolic network as the gene set in that network. We compared gene sets from each draft network with those in the corresponding manually curated model by case sensitive string comparison.

### Comparison of metabolite sets

Each metabolic network contains a set of metabolites. For those networks generated with reconstruction tools using the BIGG database (AuReMe, CarveMe and MetaDraft) we compared metabolites just by string comparison. For other reconstruction tools (Merlin, ModelSEED, Pathway Tools and RAVEN), we mapped the metabolites using MetaNetX version 3.0 [54]. As metabolite identifiers in the manually curated models contain at the end of the string a character describing the specific compartment in which the metabolite is located (for example glc_c for glucose in the cytoplasmic space) and in MetaNetX they do not, we used the following procedure to compare metabolites: For each metabolic network and for each metabolite we removed the compartment char from the metabolite identifier. Then, if the modified identifier is present in MetaNetX and if there is a synonym for that identifier in the BIGG database, we checked if some of the BIGG synonyms concatenated with the before removed compartment char match a metabolite in the manually curated model. If so, we considered that the metabolite is present in the manually curated model. Otherwise, we considered that the metabolite is not present.

### Comparison of reaction sets

Each metabolic network contains a set of reactions. Reactions sets were compared using two complementary methodologies. First, by using reaction identifier MetaNetX mapping and second, by using reaction equation comparison.

In the first approach, as a pre-processing step, we removed duplicated reactions (those reactions with the same MetaNetX identifier even if the reaction equation is different). For those networks generated with reconstruction tools using the BIGG database (AuReMe, CarveMe and MetaDraft) reactions identifiers were compared by direct case sensitive string comparison. For other reconstruction tools, MetaNetX was used to map reaction identifiers, which also were compared by string comparison.

In the second case, as a pre-processing step, we first removed duplicated reactions (those with the same equation even if they had different identifiers) and empty reactions (those with an identifier but with no reactants and products). Then, reaction equations were compared by comparing each metabolite and its stoichiometry individually. For those networks generated with reconstruction tools using the BIGG database (AuReMe, CarveMe and MetaDraft) we directly compared reaction equations. For those networks generated with reconstruction tools using a database different from BIGG (Merlin, ModelSEED, Pathway Tools and RAVEN), we first converted metabolite identifiers to BIGG by using MetaNetX version 3.0 and our own dictionary (S13 Table). Then, reaction equations were compared.

All the comparison was done in MATLAB and model handling was performed using functions from Cobra Toolbox v.3.0 [14]

### Calculation of Jaccard Distance

The Jaccard distance (*JD*) was calculated to compare reconstructions in terms of genes, reactions and metabolites. For two any sets of elements, *S_i_* and *S_j_*, the *JD* is calculated as *JD* = 1 – |*S_i_* ∩ *S_j_*|/|*S_i_* ∪ *S_j_*| We called *JD_g_, JD_r_* and *JD_m_* to the *JD* calculated in terms of genes, reactions and metabolites, respectively. Thus, *JD_g_, JD_r_* and *JD_m_* were calculated as: *JD_g_* = 1 – |*G_i_* ∩ *G_ref_*|/|*G_i_* ∪ *G_ref_*|, *G_i_* being the genes set of the generated draft network *i* and *G_ref_* being the genes set of the reference network (manually-curated model).

*JD_r_* = 1 – |*R_i_* ∩ *R_ref_*|/|*R_i_* ∪ *R_ref_*|, *R_i_* being the reactions set of the generated draft network *i* and *R_ref_* being the reactions set of the reference network (manually-curated model).

*JD_m_* = 1 – |*M_i_* ∩ *M_ref_*|/|*M_i_* ∪ *M_ref_*|, *M_i_* being the metabolites set of the generated draft network *i* and *M_ref_* being the metabolites set of the reference network (manually-curated model).

### Calculation of Ratio

The ratio (*R*) between the coverage and the percentage of additional elements was calculated to assess how similar a particular draft network was to the manually curated reconstruction. We called *R_g_, R_r_* and *R_m_* to the *R* calculated in terms of genes, reactions and metabolites, respectively. Thus, *R_g_, R_r_* and *R_m_* were calculated as:

> *R_g_* = |*G_i_* ∩ *G_ref_*|/|*G_i_* – *G_ref_*|, *G_i_* being the genes set of the generated draft network *i* and *G_ref_* being the genes set of the reference network (manually-curated model).
>
> *R_r_* = |*R_i_* ∩ *R_ref_*|/|*R_i_* – *R_ref_*|, *R_i_* being the reactions set of the generated draft network *i* and *R_j_* being the reactions set of the reference network (manually-curated model).
>
> *R_m_* = |*M_i_* ∩ *M_ref_*|/|*M_i_* – *M_ref_*|, *M_i_* being the metabolites set of the generated draft network *i* and *M_j_* being the metabolites set of the reference network (manually-curated model).

### Evaluation of performance

We created three models of Lactobacillus plantarum with CarveMe version 1.2.1 and ModelSEED version 2.4, using different media compositions for the gap-filling procedure that is carried out internally in these tools. Since the models were not able to generate biomass with the original media composition of CDM, PMM7 and PMM5 [49], we modified these mediums to ensure growth. The lack of growth was because of the presence of some compounds in the biomass equation which were not provided in the media. The modified mediums were called CMM-like, PMM7-like, PMM5-like, respectively (S2 File).

A set of 34 single-omission experiments [49] were used to evaluate the performance of the models. True positive were defined as growth *in vivo* and *in silico*; True negatives as no growth *in vivo* and *in silico*; False positives as no growth *in vivo* and growth *in silico*; False negatives as growth *in vivo* but no growth *in silico*. CDM-like media was used as a basal media for the single omission experiments. For both *in vivo* and *in silico* experiments, growth rates below 10% of the growth rate obtained in CDM-like were considered as no growth.

Metrics to evaluate the performance were calculated as follows:

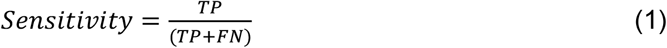

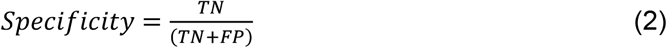

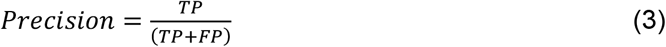

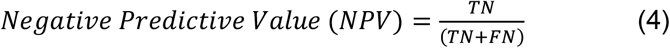

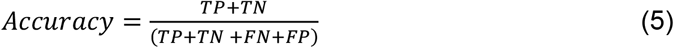

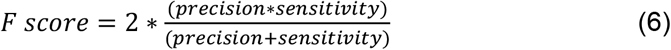

## Acknowledgements

S.N.M. acknowledges the financial support from CONICYT Becas Chile #72180373 and Chr. Hansen

## Authors contributions

### Conceptualization

Sebastián N. Mendoza, Brett G. Olivier, Douwe Molenaar, Bas Teusink

### Formal Analysis

Sebastián N. Mendoza

### Funding Adquisition

Sebastián N. Mendoza, Bas Teusink

### Methodology

Sebastián N. Mendoza, Brett Olivier, Douwe Molenaar, Bas Teusink

### Resources

Bas Teusink

### Software

Sebastián N. Mendoza

### Supervision

Bas Teusink

### Writing - original draft

Sebastián N. Mendoza

### Writing - review & editing

Sebastián N. Mendoza, Brett G. Olivier, Douwe Molenaar, Bas Teusink

## Availability

All the reconstructions used as well as the MATLAB functions to generate the models (when possible) and to compared them are available at https://github.com/SystemsBioinformatics/pub-data/reconstruction-tools-assessment.

## Abbreviations

FBA: Flux Balance Analysis
GSMM: Genome-scale metabolic model
JD: Jaccard Distance
LAB: lactic acid bacterium.
R: Ratio between the coverage and the percentage of additional elements.
SBML: Systems Biology Markup Language

## Supporting information captions

**S1 Fig. Performance of models made with CarveMe and ModelSEED for *Lactobacillus plantarum* when different media compositions were provided for the internal gap-filling performed in these tools.** Networks gap-filled with CDM-like got an accuracy lower but close to the obtained with the manually-curated model. For both tools, when PMM7-like or PMM5-like was used, the accuracy score increased due to the decrease in false negative results. For both tools, the lines corresponding to PMM7-like are not visible because they overlap with the ones of PMM5-like.

**S1 Table. List of genome-scale metabolic reconstruction tools and databases.**

**S2 Table. Specification of scores in each feature evaluated.**

**S3 Table. Detailed evaluation of the reconstruction tools**

**S4 Table. Calculated Jaccard Distance between genes, metabolites and reactions sets in draft networks of *L. plantarum* and those in iLP728.**

**S5 Table. Calculated Jaccard Distance between genes, metabolites and reactions sets in draft networks of *B. pertussis* and those in iBP1870.**

**S6 Table. Similarity of draft reconstructions of *L. plantarum* to iLP728 in terms of reactions, metabolites and genes.**

**S7 Table. Similarity of draft reconstructions of *B. pertussis* to iBP1870 in terms of reactions, metabolites and genes.**

**S8 Table. Metabolic processes associated to the 92 genes of iLP728 which were recovered in all the draft networks of Lactobacillus plantarum**

**S9 Table. Metabolic processes associated to the 121 genes of iBP1870 which were recovered in all the draft networks of Bordetella pertussis**

**S10 Table. Reaction coverage for draft reconstructions of *L. plantarum* using different approaches.**

**S11 Table. Reaction coverage for draft reconstructions of *B. pertussis* using different approaches.**

**S12 Table. New unique reaction synonyms pairs that were automatically discovered for both species with the reaction equation comparison approach.**

**S13 Table. Compound synonyms pairs that were semi-automatically discovered.**

**S14 Table. Additional reaction synonyms discovered.**

**S15 Table. Metabolic processes associated to the 71 reactions of iLP728 which were recovered in all the draft networks of Lactobacillus plantarum.**

**S16 Table. Metabolic processes associated to the 91 reactions of iBP1870 which were recovered in all the draft networks of Bordetella pertussis.**

**S167 Table. Classification of reactions in iLP728 which were not recovered by any draft network of *L. plantarum*.**

**S18 Table. Classification of reactions in iBP1870 which were not recovered by any draft network of *B. pertussis*.**

**S19 Table. Metabolic processes associated to the 208 metabolites of iLP728 which were recovered in all the draft networks of Lactobacillus plantarum.**

**S20 Table. Metabolic processes associated to the 266 metabolites of iBP1870 which were recovered in all the draft networks of Bordetella pertussis.**

**S21 Table. Metabolic processes associated to the 9 metabolites which were recovered in all the draft networks of *L. plantarum* and were not in iLP728.**

**S22 Table. Metabolic processes associated to the 13 metabolites which were recovered in all the draft networks of Bordetella pertussis and were not in iBP1870.**

**S23 Table. Topological analysis for draft recontructions of *L. plantarum*.**

**S24 Table. Topological analysis for draft recontructions of *B. pertussis*.**

**S1 File. Specific input and parameters set to obtain each of the reconstructions studied.**

**S2 File. Composition of CDM-like, PMM7-like and PMM5-like mediums.**

